# Integrin-mediated adhesion in the unicellular holozoan *Capsaspora owczarzaki*

**DOI:** 10.1101/2020.02.27.967653

**Authors:** Helena Parra-Acero, Matija Harcet, Núria Sánchez-Pons, Elena Casacuberta, Nicholas H. Brown, Omaya Dudin, Iñaki Ruiz-Trillo

## Abstract

In animals, cell-matrix adhesion is mainly mediated by integrins and their associated proteins. Comparative genomic analyses have shown that core components of the integrin adhesome pre-date the emergence of animals, however, whether it mediates cell adhesion in non-metazoan taxa remains unknown. Here, we investigate cell-substrate adhesion in *Capsaspora owczarzaki*, the closest unicellular relative of animals with the most complete integrin adhesome. Using an adhesion assay, we show that *C. owczarzaki* adheres to surfaces using actin-dependent filopodia. We show that integrin β2 and its associated protein vinculin localise as distinct patches in the filopodia. We also demonstrate that substrate adhesion and integrin localisation are enhanced by the ligand fibronectin. Finally, using a specific antibody for Integrin β2, we inhibited cell adhesion to a fibronectin-coated surface. Our results show that adhesion to the substrate in *C. owczarzaki* is mediated by integrins. This suggests that integrin-mediated adhesion pre-dates the emergence of animals.

## Introduction

In animals, cell-matrix adhesions are essential for cell migration, tissue organisation, and differentiation, which have central roles in embryonic development (Berrier and Yamada, 2007; Bulgakova et al., 2012; Hynes, 1999; Hynes and Zhao, 2000; Maartens and Brown, 2015; Winograd-Katz et al., 2014). Integrins are the major cell surface adhesions receptors mediating cell-matrix adhesion in animals. They are heterodimeric transmembrane proteins that bind extracellular matrix (ECM) molecules on one side and connect to the actin cytoskeleton on the other (Hynes, 2002). Various combinations of α and β integrin subunits bind to different ECM ligands, such as fibronectin, collagen and laminin, with different affinities (Barczyk et al., 2010; Campbell and Humphries, 2011; Humphries, 2006). The connection to the cytoskeleton is mediated by integrin-associated proteins (IAPs) (Green and Brown, 2019; Zaidel-Bar et al., 2007), among which, talin and vinculin are known to interact directly with actin filaments and actin nucleators such as Arp2/3 complex (Bays and DeMali, 2017; Klapholz and Brown, 2017). Given the importance of integrin-mediated cell-matrix adhesion in development and homeostasis of multicellular animals, it is of interest to discover when and how this machinery arose during evolution.

Comparative genomic analyses have recently shown that core elements of the integrin adhesome pre-date the origins of Metazoa (Brown et al., 2013; de Mendoza et al., 2015; Hehenberger et al., 2017; Sebé-Pedrós et al., 2010). However, whether integrin adhesome plays a role in cell-matrix adhesion in non-metazoans, or has some other function, has yet to be investigated. Among holozoans — a clade that includes animals and their closest unicellular relatives (Lang et al., 2002) (Figure 1A) — the filasterean *Capsaspora owczarzaki* contains the greatest number of orthologs of integrin adhesome components amongs unicellular organisms examined to date (Sebé-Pedrós et al., 2010; our unpublished observations). Its genome encodes four integrin α and four integrin β subunits as well as most of other essential components of the integrin adhesion complex, including focal adhesion kinase and tensin, which are absent from other unicellular organisms examined to date (Sebé-Pedrós et al., 2010; Sebé-Pedrós and Ruiz-Trillo, 2010; our unpublished observations). The life-cycle of *Capsaspora owczarzaki (*hereafter, *Capsaspora*) includes at least three distinct life stages; adherent, cystic and aggregative (Sebé-Pedrós et al., 2013b). Transcriptomic profiling showed that main components of the integrin adhesome, as well as proteins containing fibronectin and laminin domains, are expressed throughout the life-cycle and up-regulated during the aggregative stage (Sebé-Pedrós et al., 2013b). These results suggest that the integrin adhesome might play a role in mediating interactions between cells to promote aggregation, possibly via secreted ligands. This role, and possible roles in mediating movement over a substrate have yet to be tested.

**Figure 1.**
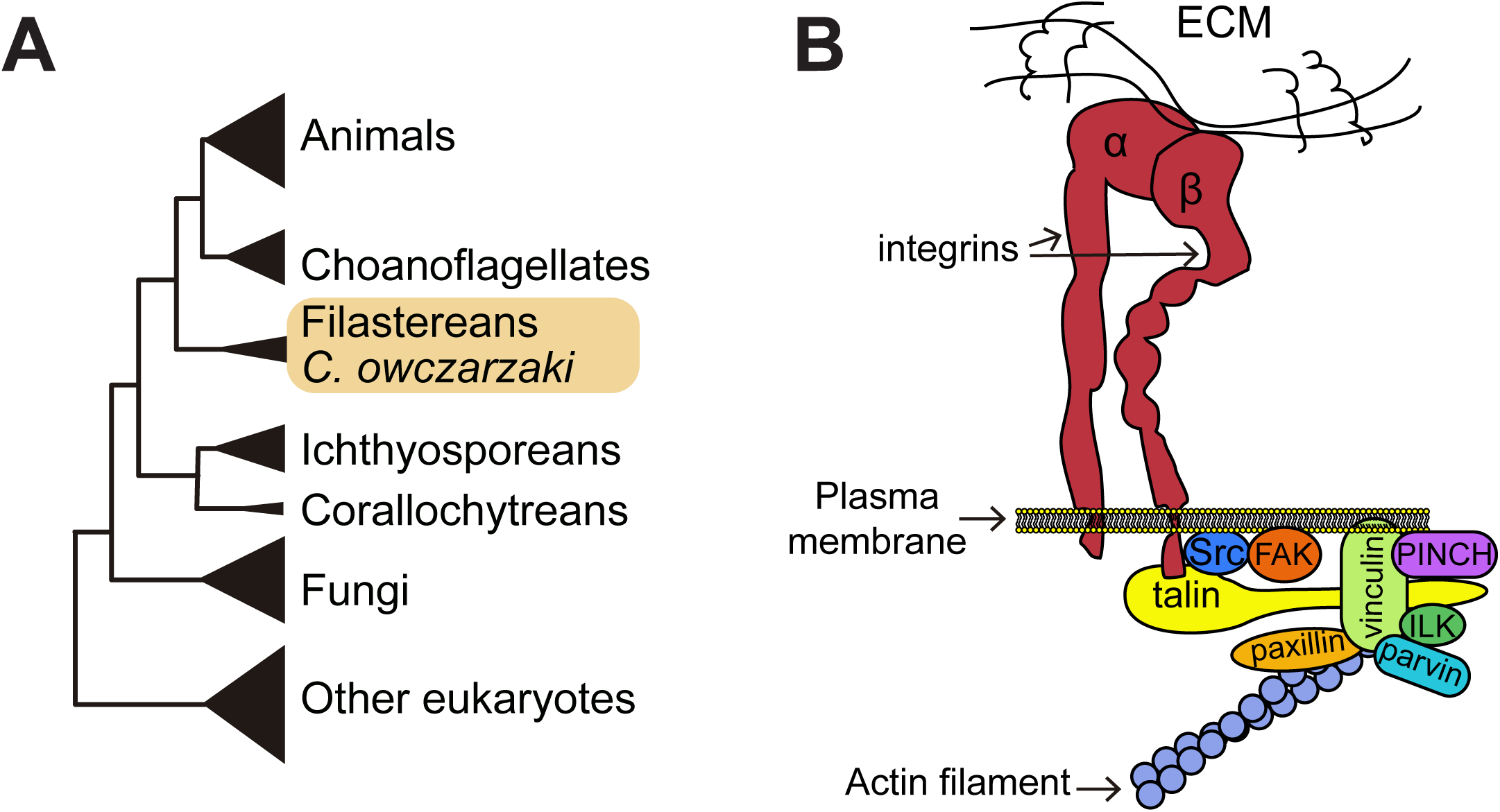
Phylogenetic position of *Capsaspora owczarzaki* and illustrative summary of the animal integrin adhesome. (A) Phylogenetic position of *Capsaspora owczarzaki* in the tree of life (adapted from Torruella et al. 2015). (B) Schematic drawing of the integrin adhesion complex in animals (adapted from Sebé-Pedrós et al. 2010).

In this work, we used adhesion assays, immunolabelling and chemical inhibition to investigate cell adhesion in the unicellular relative of animals *Capsaspora*. We show that *Capsaspora* can adhere to a fibronectin coated surface via actin-dependent filopodia. Using immunolabelling, we show that integrin β2 and vinculin localise as distinct patches along the filopodia. Such localisation is enhanced on a fibronectin-coated surface. Finally, adhesion to fibronectin coated surfaces is inhibited by the integrin β2 antibody. Our results indicate that orthologs of the animal integrin adhesome are involved in substrate adhesion in *Capsaspora*. This suggests that the role of these proteins in cell-matrix adhesion was established before the emergence of animals.

## Results

Exponentially growing *Capsaspora* are thought to be in an adherent life-stage consisting of single cells adhering to the substrate (Sebé-Pedrós et al., 2013b). However, the capacity of *Capsaspora* to adhere to a surface has never been systematically investigated. To achieve this, we modified previously published adhesion assays (Busk et al., 1992; Pierschbacher and Ruoslahti, 1984) (Figure 2A). Briefly, we seeded exponentially growing cells into an untreated multi-well plate and let them sit for 2.5 hours. We then discarded the cells remaining in suspension and fixed the remaining cells attached to the bottom of the well. After several washes, we indirectly measured the number of cells that remained adhered to the surface by quantifying DAPI staining of nucleic acids (Figure 2A, Materials and Methods). Using this assay, we observed that fluorescence intensity increased with cell concentration (Figure 2B, 2C). This assay is most sensitive between 1.6×10^6^ cells/mL and 6.6 ×10^6^ cells/mL (Figure 2C), so we seeded 3.3 ×10^6^ cell/mL for all future experiments to allow us to measure with confidence any increase or decrease in cell adhesion. Our results show that *Capsaspora* can adhere to surfaces and the adherent cells are quantitatively measured with our assay.

**Figure 2.**
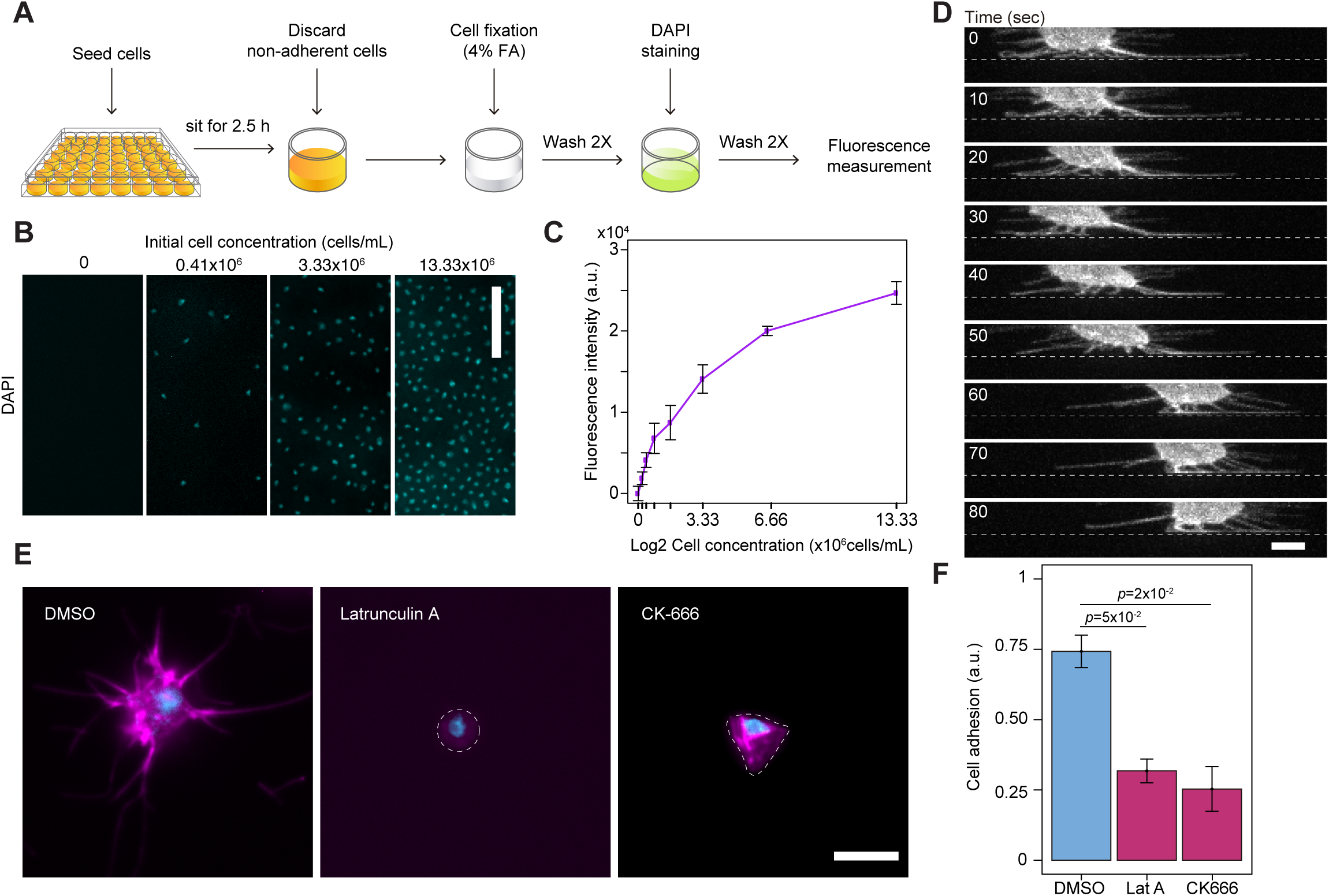
*Capsaspora owczarzaki* adheres to surfaces using actin-dependent filopodia. (A) Protocol for measuring cell adhesion in *Capsaspora*. (B) DAPI-stained cells remaining attached to the surface following the adhesion assay. Number of seeded cells at the beginning of the assay is indicated. Scale bar = 50µm. (C) Fluorescence intensity measures of an adhesion assay performed with increasing concentration of *Capsaspora* cells seeded on untreated plates. (D) Time-lapse of an adherent *Capsaspora* cell transfected with a membrane label (CoNMM:mcherry). Images were taken as a Z-stack every 10 sec. A maximum intensity projection of an orthogonal view is represented. Punctate line marks the substrate. Scale bar = 5µm. (E) Phalloidin (magenta) and DAPI (cyan) staining of floating cells from an adhesion assay treated with 0.05mM LatA or 0.1 mM CK666, and its corresponding DMSO control. Scale bar =5µm. (F) Adhesion assay in presence of drugs. Data represent the ratio of signal from adherent cells treated with Lat A, CK666 or DMSO relative to the signal of untreated cells. Bars represent mean ± s.e.m. (n=3) of three independent experiments. As both DMSO controls show similar effect, only one is shown for simplicity. *p-values* from a paired t-test are shown.

Initial descriptions of *Capsaspora* showed that it has dynamic filopodia (Stibbs et al., 1979), and a recent study revealed that these filopodia contact the substrate and appear to hold the cell body above the surface (Parra-Acero et al., 2018). We examined filopodia behaviour in live cells, by performing time-lapse microscopy of adherent *Capsaspora* cells transfected with a membrane marker (NMM-mCherry) (Parra-Acero et al., 2018). We observed that filopodia not only hold the cell over the surface but appear to actively participate in its motility (Figure 2D). This suggests that filopodia are the major interactor with the surface, and thus potentially crucial for cell-surface adhesion. In animals, filopodia formation and maintenance relies extensively on the actin cytoskeleton (Mattila and Lappalainen, 2008; Mellor, 2010). Several actin regulators involved in filopodia formation, including Arp2/3 complex, a major actin nucleator in animals (Mellor, 2010; Rottner et al., 2017) are found in *Capsaspora* (Sebé-Pedrós et al., 2013a). We thus used Latrunculin A (LatA), a general inhibitor of actin polymerisation, and Arp2/3-specific inhibitor CK-666 to assess the role of filopodia in adhesion (Burke et al., 2014; Yarmola et al., 2000). We observed that LatA treated cells were round and lacked both filopodia and actin filaments (Figure 2E). CK-666 treated cells did not lack actin filaments and maintained an elongated cell shape, but completely lacked filopodia (Figure 2E). Adhesion assays of LatA and CK-666 treated cells showed a substantial decrease in adhesion (Figure 2F), suggesting that cell-surface adhesion in *Capsaspora* is mediated by Arp2/3-dependent filopodia.

In animals, cell-matrix adhesions are formed by the interaction of integrins and cytoplasmic integrin-associated proteins, such as vinculin, at discrete sites in the cell (Kanchanawong et al., 2010; Zamir and Geiger, 2001). To address whether conserved components of the integrin adhesome in *Capsaspora* are also involved in cell adhesion, we developed antibodies for immuno-localisation. We successfully obtained functional antibodies against both the cysteine-rich region of integrin β2 (Anti-β2E3), and vinculin (Figure 3-figure supplement 1). The integrin β2 is the mostly highly expressed of the four *Capsaspora* β subunits, contributing 68% and 73% of the beta subunit mRNA (FPKM) at filopodia and aggregative stages (Sebé-Pedrós et al., 2013b). We observed that both antibodies stain the cell body and multiple distinct patches along filopodia (Figure 3A,3B). Co-immunostaining showed a partial overlap, some patches contain both, whereas others have just one of the proteins (Figure 3C). Taken together, these results suggest that the integrin and vinculin patches may represent anchorage sites of adhesion.

**Figure 3.**
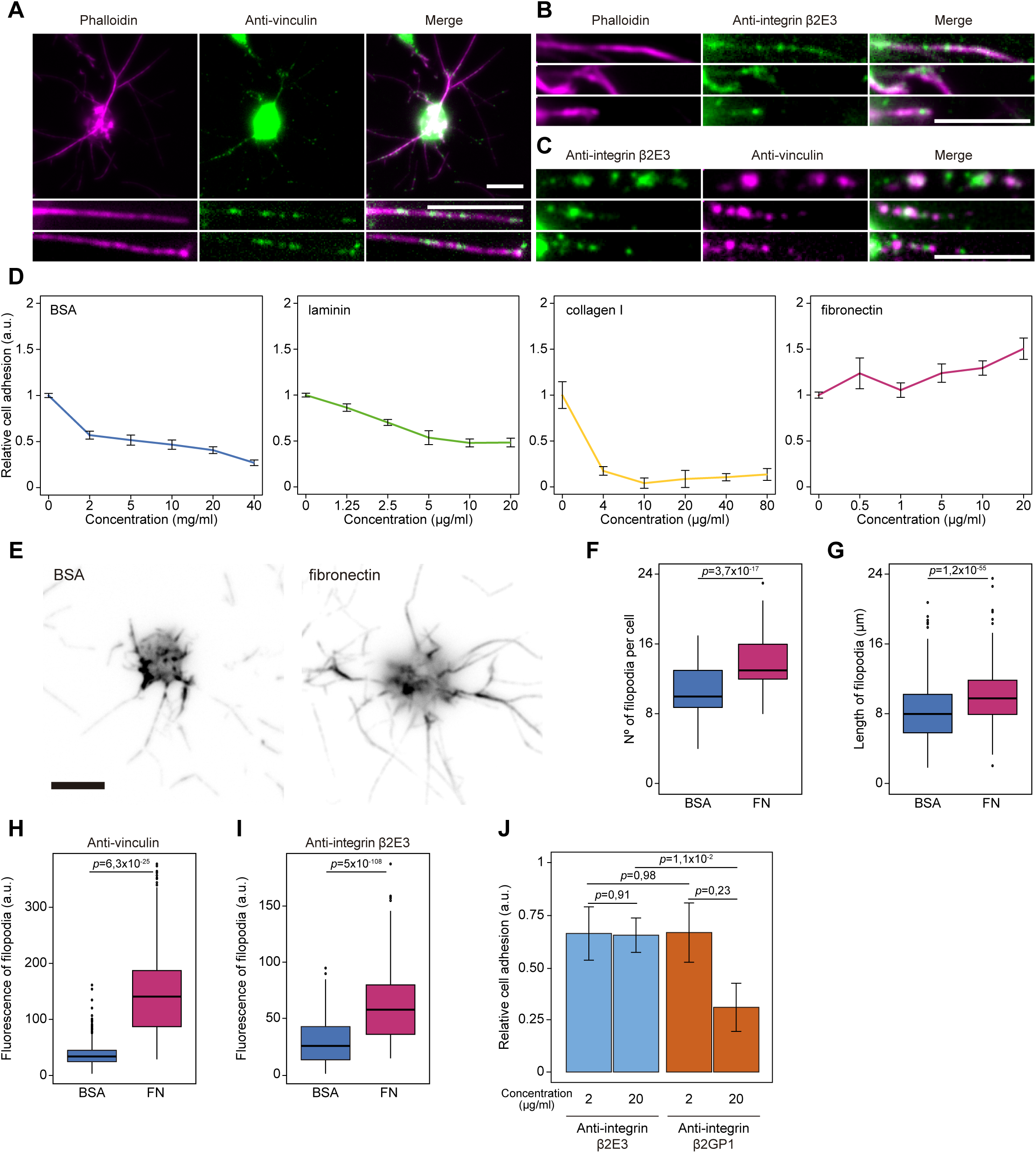
*Capsaspora owczarzaki* undergoes integrin-mediated cell adhesion on fibronectin-coated surfaces. (A) Immunostaining of adherent and phalloidin-stained *Capsaspora* cells (magenta) with anti-vinculin antibody (green). Distinct patches of vinculin are observed in the filopodia. Scale bar =5µm. (B) Immunostaining of adherent and phalloidin-stained *Capsaspora* cells (magenta) with antibody for integrin β2 subunit, anti-β2E3 (green). Distinct patches of integrin β2 are observed in the filopodia. Scale bar =5µm. (C) Co-immunostaining of vinculin (magenta) with integrin β2 (green) shows partial co-localisation. Scale bar =5µm. (D) Cell adhesion measurements on surfaces treated with increasing concentrations of bovine serum albumin (BSA), laminin, collagen I and fibronectin. BSA was included as a control. Relative cell adhesion was set as the ratio between the signal obtained between ligand-coated and uncoated surfaces. Data represents the mean ± s.e.m of 4 technical replicates. (E) Phalloidin-stained cells seeded on either fibronectin or BSA treated coverslips. Scale bar= 5µm. (F) Number of filopodia per cell seeded on a either fibronectin or BSA treated coverslips (n=164). *p-value* from a Mann-Whitney test is shown. (G) Filopodia length of cells seeded on a either fibronectin or BSA treated coverslips (n=500). *p-value* from a Mann-Whitney test is shown. (H) Vinculin fluorescence intensity of filopodia (n>100), from cells seeded on a either fibronectin or BSA treated coverslips. *p-value* from a Mann-Whitney test is shown. (I) Integrin β2 fluorescence intensity of filopodia (n>100), from cells seeded on a either fibronectin or BSA treated coverslips. *p-value* from a Mann-Whitney test is shown. (J) Cell adhesion of *Capsaspora* on fibronectin-coated surfaces in presence of different concentrations of anti-β2E3 and anti-β2GP1 antibodies. Relative cell adhesion was set as the ratio of signal obtained between cells treated with integrin antibodies and control (buffer). Data represents the mean ± s.e.m. (n=3) of 3 replicates, *p-value* from a paired t-test is shown.

To further investigate that, we first assessed whether *Capsaspora* adheres preferentially to any of the well-known integrin ECM ligands in animals; namely fibronectin, laminin and collagen 1. Coating the plastic substrate with Bovine Serum Albumin (BSA), laminin or collagen reduced the adhesion of Capsaspora (Figure 3D, Figure 3-figure supplement 2). This suggests that a protein secreted by *Capsaspora* sticks to the plastic substrate and provides a ligand for adhesion, and the binding of this protein to the plastic substrate is reduced by prior coating with BSA or laminin or collagen. In contrast, fibronectin does not reduce adhesion, and in fact increases cell adhesion with increasing concentration (Figure 3D, Figure 3-figure supplement 2). This indicates that *Capsaspora* can use fibronectin as a substrate for cell adhesion. We then assessed whether adhesion to fibronectin affects filopodia morphology or localisation of integrin β2 and vinculin to the filopodia. The number of filopodia, as well as their length increased (Figure 3E-3G). The overall fluorescence intensity of integrin β2 and vinculin staining within filopodia increased (Figure 3H,3I), and the overall distribution remained similar, with a patchy appearance (data not shown). These results suggest that Fibronectin promotes cell adhesion by increasing the number and length of filopodia as well as by promoting recruitment of integrin and vinculin to the filopodia. *Capsaspora* does not have an ortholog of fibronectin, but does have proteins with fibronectin domains (Sebé-Pedrós et al., 2013b; Suga et al., 2013) as well as secreted proteins with RGD sequences, which are candidates for endogenous ligands.

Finally, to directly assess the role of the β2 integrin cell-substrate adhesion, we generated a new antibody to the β2 integrin subunit, this time targeting the ligand-binding β-I domain (anti-β2GP1) (Figure 3-figure supplement 1). Pre-incubation of this cells with this antibody prior to plating substantially reduced adhesion to fibronectin-treated surfaces, in contrast to the antibody against the cysteine-rich stalk, which did not impair adhesion (Figure 3J).

Taken together, these results show that the major integrin β subunit in *Capsaspora* makes a substantial contribution to substrate adhesion. The localisation of integrin β2 and vinculin in discrete patches within the filopodia suggests that these structures contribute to cell-substrate adhesion, which is an unexpected role for integrin adhesion within filopodia. In animal cells filopodia are more generally associated with dynamic exploration of the substrate rather than forming stable adherent contacts. Our findings provide further support to the view that integrin-mediated cell adhesion evolved before animals emerged. Further investigation of integrin-mediated adhesion in *Capsaspora*, particularly during the aggregative stage, by examining its ligands and intracellular binding partners, will shed more light into the origins of the integrin-mediated adhesion and its role in the evolution of animal multicellularity.

## Author contributions

H.P.-A, M.H., O.D. and I.R-T., designed the study and set up the methodology. H.P.-A and O.D. performed the adhesion assays, the microscopy and the chemical inhibition experiments. N.S.-P., H.P.-A and M.H. designed and validated the antibodies. N.H.B. and E.C. assisted in the design of the project and analysis of results. I.R.-T. and O.D. provided supervision. H.P.-A and O.D. wrote the original draft. All authors reviewed and edited the manuscript.

## Acknowledgments

We thank Meritxell Antó for technical support. We thank Núria Ros-Rocher for assistance with transfection of the organism. We acknowledge the Biomolecular Screening and Protein Technologies Unit from CRG, (Barcelona,Spain) for assistance with antibody production; as well as the CRG/UPF Proteomics Unit for proteomics analysis. We also thank CRG Advanced Light Microscopy Unit for support with confocal imaging. We also thank the many other people who have worked in our lab and who have provided support and feedback over the years.

This work was funded by European Research Council Consolidator Grant (ERC-2012-Co -616960) to I.R.-T.; O.D. was supported by a Swiss National Science Foundation Early PostDoc Mobility fellowship (P2LAP3_171815) and a Marie Sklodowska-Curie individual fellowship (MSCA-IF 746044). M.H. received funding from the People Programme (Marie Curie Actions) of the European Union’s Seventh Framework Programme FP7/2007-2013/under REA grant agreement no. 330925. N.H.B.’s work on this project was supported by a Royal Society International Exchange Grant (IE141189). The CRG/UPF Proteomics Unit is part of the of Proteored, PRB3 and is supported by grant PT17/0019, of the PE I+D+i 2013-2016, funded by ISCIII and ERDF

## Materials and Methods

### Cell strain and growth conditions

*Capsaspora owczarzaki* cell cultures (strain ATCC®30864) were grown axenically at 23°C in ATCC medium 1034 (modified PYNFH medium). 25 cm^2^ and 75 cm^2^ flasks were used for culture (Falcon®, #353108 and #353136 respectively). Adherent cells were obtained by growing cells for few days and before floating cells appeared.

### Custom polyclonal antibodies design

β2E3-antigen was designed as a region in the stalk of the extracellular domain of the integrin β2 (CAOG_05058, aminoacid region 733-835). The sequence (5’ CAATGCGTCTGCGATGCTTTGCACGCCGGCCCTGCTTGCGGCTGCGTCAAGGGTGTCTGCCCCTCTGTTGGCGGCGTTCGCTGCAATGGCGGTGATTGCGACCCAATCTGCGGTATCTGCACTTGCCCGCCTGGCAAGACTGGACCTGCGTGCGACTGCGATACTGTTGCTCACCCGTGCCCGACTGGCAACTCCACCTCTGGCGTTGTGCTTCCCTGCTCTGGCCAAGGCACGTGCCTGCAGTCGTCTGCCACTCAGTGCGGCATCTGCTTGTGCAACCGTGATCCGCTGACTGGCACGCCGCTGTAC 3’) corresponding to that region was amplified using the following primers: forward (5’ CTGAATTCCAATGCGTCTGCGATGC 3’) and reverse (5’ AGACTCGAGGTACAGCGGCGTG 3’) and cloned into plasmid pGEx4T1 using *EcoR*I and *Xho*I restriction enzymes. The polypeptide was produced in *E.coli* by the Biomolecular Screening and Protein Technologies (BSM&PT) Unit of CRG, Barcelona; and its corresponding polyclonal antibody was produced in guinea-pig by TebuBio and affinity purified by BSM&PT, CRG. β2GP1-antigen is a polypeptide corresponding to a region in the β-I domain of integrin β2 (aminoacid region 100-403). The vinculin-antigen is a polypeptide corresponding to the C-terminus region of one of the vinculin homologs (CAOG_05123, amino acid region 335-834). Antigens for both β2GP1 and vinculin, and their corresponding polyclonal antibodies (in guinea-pig and rat respectively) were designed and produced by Genecust. Antibody stock concentration were: anti-β2E3 1.8 mg/mL, anti-vinculin and anti-β2GP1 1mg/mL.

### Protein extraction

Protein extraction was performed as follows: adherent and exponentially growing cells were scraped and pelleted at 5000 x g. Cells were then washed once with PBS 1X and kept at either -20°C or - 80°C. Cells were resuspended in lysis buffer (Tris-HCl 50mM pH 8.8, NaCl 150mM, SDS 0.1%, EDTA 5mM, EGTA 1mM, NP-40 1%, MgCl2 1mM, CaCl2 1mM, DTT 1mM, PMSF 0.5mM, half cOmplete (4693159001, Sigma-Aldrich) for 3mL buffer) and kept on ice for 10 min before being sonicated (amplitude 10%, 3 pulses of 15s, 45s between pulses). To obtain the final protein extract was obtained after centrifugation for 30 min at 20,000 x g at 4°C and the supernatant was kept as the soluble fraction.

### Western Blot

For SDS-PAGE, 1-5 μg of purified antigen or 20 μg protein extract were loaded in a precast 4-20% acrylamide gel (BioRad 456-1094). Gels were run for the first 20-30 min at 40V, later at 80-100V. Proteins and antigens were wet-transferred in TGS 1X (Sigma T7777-1L) with methanol 20% to a nitrocellulose membrane (Amersham 10600004) at 20-30 V 4°C overnight. After transfer, membranes were incubated with blocking solution (Tween 0.1% and 5% no-fat powder milk in PBS 1X) at room temperature (RT) for 1 hour. Membranes with protein extract were quickly washed with 0.1% Tween in PBS 1X (PBS-T hereafter), and incubated with primary antibodies at 4°C overnight in the following conditions: anti-IntB2E3 3.6 µg/mL in PBS-T with 1% milk, anti-IntB2GP1 and anti-vinculin 2µg/mL in PBS-T. Membranes with antigens were incubated with primary antibody 2h at RT in the following conditions: anti-β2E3 1.8µg/mL in PBS-T with 1% milk (no-fat, powder), anti-β2GP1 and anti-vinculin 1µg/mL in PBS-T. After incubation, membranes were washed 4 times with PBS-T and incubated with secondary antibodies: anti-guinea pig HRP (Invitrogen, #61-4620), and anti-rat HRP (Abcam, ab7097) at 0.1µg/mL in PBS-T for 3 h (2 h for antigens) at RT. After incubation with secondary antibody, membranes were washed again with PBS-T. Detection was performed by chemiluminescence (SuperSignalTM West Pico Chemiluminescent Substrate 34078 Thermofisher).

### Immunoprecipitation and Mass-spectrometry

For immunoprecipitation (IP), protein extraction was performed as previously described, in lysis buffer without DTT or SDS, and 1% Triton x100 instead of NP-40. 1 mg of protein extract was used for IP with vinculin antibody and 1.5mg for IP with the β2E3 and β2GP1 antibodies. For IP with guinea-pig antibody protein A-conjugated magnetic beads (Merck Millipore, LSKMAGA10) were used. For IP with rat antibody, protein G-magnetic beads (Merck Millipore, LSKMAGG10) were used instead. The lysate was pre-cleared by incubating with 50µL of beads per condition overnight at 4°C. Beads were previously washed twice with IP-buffer (PBS-Tween 0.1%). Additionally, 50uL beads per antibody were washed with IP-buffer and blocked with blocking solution (1%BSA in PBS-Tween 0.1%) for 1h at room temperature. Blocked beads were mixed with a dilution of each antibody in IP-buffer (9µg/mL of anti-β2E3 and 5µg/mL of anti-β2GP1 and anti-vinculin) and incubated 4h at 4°C, then washed three times with IP-buffer. Pre-cleared lysate was mixed with antibody-bound beads and incubated overnight at 4°C. Beads were then washed with IP-buffer three times and kept without liquid at -20°C. Negative controls with guinea-pig IgG (MBL PM067) and rat IgG (Sigma, I8015) (5µg/mL) were made following the same procedure. “In-beads” digestion and Mass Spectrometry was performed in the CRG/UPF Proteomics Unit in Barcelona, Spain.

### Membrane labelling

*Capsaspora* was transfected in its adherent stage according to a calcium phosphate precipitation method (Parra-Acero et al. 2018) with the *CoNMM:mcherry* construct, which directs mCherry to the cell membrane, (Parra-Acero et al 2018). Transfected cells were seeded in a µ-Slide 4-well glass-bottom dish (Ibidi, #80427) and grown overnight at 23°C prior to imaging.

### Adhesion assay

Adherent cells were obtained as mentioned above and then scraped and harvested by centrifugation at 5000 x g before being re-suspended in growth medium without fetal bovine serum (FBS) (FBS presence showed heterogeneous results, data not shown). Cells were then seeded in untreated 48-well plate (32048, SPL Life Sciences) at a concentration of 3.3 ×10^6^ cells/ml. Cells were left to sit and adhere for 2.5 hours at 23°C. The medium containing floating cells, was then gently removed and adherent cells were fixed with 4% formaldehyde for 10 min at room temperature (RT). The wells containing fixed cells were washed once with 1X PBS and stained with DAPI (50 μg/ml) for (Roche, 10236276001) 10 min at RT and protected from light. Finally, wells were washed 3 times with 1X PBS followed by immediate fluorescence measurement with a plate reader (TECAN infinite 200). Fluorescence of DAPI (350nm excitation /470nm emission) was read using the i-control™ Microplate Reader Software. Detection was performed from the top, set to 5×5 reads per well (circle filled) and gain was set to “optimal” automatically by the software for each independent experiment. Unless pointed otherwise, four technical replicates were performed for each condition per experiment. During analysis, background signal was removed. For easier visualisation of the results, fluorescence measurements were normalised to the control of seeded cells in untreated wells. Equal number of cells adhering in uncoated and coated wells or treated and untreated cells would set a value equal to 1.

For assessing the role of actin in cell-surface adhesion, both Latrunculin A (LatA) (Sigma-Aldrich L5163-100UG) and CK-666 (SML0006-5MG) were used as pharmacological inhibitors. Stock solutions was 20mM for LatA and 10mM for CK-666 in DMSO. Re-suspended cells were incubated for 10 minutes with both pharmacological inhibitors at a final concentration of 50 μM of LatA and 100 μM of CK666, and its corresponding DMSO amount as control. Then, cells were plated and the treatment continued during the sitting time. Adhesion assay was then preformed as mentioned above.

For ECM-ligand affinity measurements, untreated multi-well plates were coated with different concentration of fibronectin (Sigma F1141-2MG), laminin (Sigma L2020-1MG), collagen I (Sigma C3867-1VL) or BSA (Sigma A3294-10G). BSA was included as control as it acts as a blocking agent of unspecific adhesion sites (Bourdon and Ruoslahti, 1989; Busk et al., 1992; Weinreb et al., 2004; Yamada and Kennedy, 1984). For all proteins, solutions of the desired concentration were prepared by diluting the stock solutions in PBS 1X. Fibronectin-coated plates were incubated overnight at 4°C and washed once with PBS 1X before plating the cells. Laminin-coated plates were incubated 2 hours at 37°C and washed 3 times with PBS 1X before plating the cells. Collagen I-coated and BSA-coated plates were incubated 1 hour at room temperature (18°C culture room) and washed once with PBS1X before plating the cells.

For assessing role of integrin inhibition using competition with anti-integrin antibodies, anti-β2E3 and anti-β2GP1 were diluted in medium without FBS and mixed with re-suspended cells, to reach a final concentration of 2 and 20μg/mL of each antibody. Then, cells mixed with antibodies were seeded on a fibronectin-coated plate (20μg/ml). Negative controls were prepared by replacing antibodies with corresponding buffer.

### Immunolocalisation of adhesome proteins

Coverslips used for immunostaining were previously coated with either BSA 4% or Fibronectin at 20 μg/mL and incubated overnight at 4°C prior to being washed once with PBS1X. Cells were scraped from a confluent culture and resuspended in growth medium without FBS before being added over the coated coverslips inside a 6 well/plate and incubated at 23°C for 2.5 hours. Excess liquid was then removed before fixing the cells with 4% formaldehyde (in PBS1X) for 10 min at RT. Cells were then washed with PBS 1X and incubated 2h at RT with primary antibodies diluted in blocking solution (1% BSA, 0.1% Triton in PBS 1X) at the following concentrations: anti-β2E3 (9μg/mL), anti-vinculin (10 μg/mL). Cells were washed with blocking solution (once quick, twice for 10 min) and incubated 1 hour at RT with secondary antibodies diluted in blocking solution as follows: anti-guinea pig Dylight 488 (Thermo Scientific, SA5-10094) (0.5 μg/mL) and anti-rat Alexa 488 (Invitrogen, A-11006) (1 μg/mL). Cells were washed with PBS 1X and incubated with phalloidin-Texas red (Invitrogen T7471) (1:100 from stock, equals 2 units/mL) 15 min in darkness. They were mounted on glass slides with ProlonGold (Invitrogen, P36930). Negative controls lacking primary antibody were performed in the same conditions and used to adjust the imaging settings.

For co-immunostaining, cells were incubated at the same time with anti-β2E3 (9μg/mL) and anti-vinculin (10 μg/mL). Secondary antibodies used were anti-guinea pig Dylight 488 (at 0.5 μg/mL) and anti-rat Texas red (1μg/mL). Cells were finally stained with phalloidin Alexa 350 (Invitrogen A22281) (2 units/mL) for 15 min at RT.

For actin localisation, cells treated with Lat A (0.05mM) and CK666 (0.1 mM) and DMSO were fixed adding 4% formaldehyde directly in the medium for 10 min. Cells were then pelleted by repeated centrifugations of 5 min at 3000 g, where the liquid was carefully removed. Cells were washed with PBS 1X and stained with Phalloidin Alexa 546 (Invitrogen A22283) (2 units/mL) and DAPI (10μg/mL) for 15min in darkness at RT.

### Microscopy

Images of fixed cells were obtained using a Zeiss Axio Observer Z.1 Epifluorescence inverted microscope equipped with Colibri LED illumination system and an Axiocam 503 mono camera using a Plan-Apochromat 63X/1.4 oil objective. Confocal microscopy of the transfected cell was obtained using an Andor Revolution XD imaging system equipped with a Olympus inverted microscope, using a 60x oil objective, a spinning disk scanhead (Yokogawa CSU-X1) and an Andor Ixon 897E Dual Mode EM-CCD camera. Images were edited using Fiji Imaging Software (version 2.0.0-rc-44/1.50e) (Schindelin et al. 2012).

### Image analysis

Image analysis was done using ImageJ software (version 1.52) (Schneider et al., 2012). For measurements of number and length of filopodia we used the ObjectJ plugin in ImageJ and measured the filopodia from the surface plane of fixed cells images. Only filopodia connected to the cell body were counted (filopodia that were broken during the staining procedure were ignored). The mean fluorescence intensity of integrin and vinculin antibody staining was measured along manually selected filopodia (over 100 filopodia). A rectangular selection covering the width and the length of the filopodia (excluding the cell body) was used to measure the mean fluorescence intensity. Fluorescence intensity was corrected for background fluorescence and normalised to the length of the corresponding filopodium.

### Statistical Analysis

Results from drug effect and antibody on adhesion assay are shown as mean ± standard error of the mean (s.e.m) as from 3 independent experiments. The significance of difference in the mean was tested using the parametric t-test for paired samples. Results from measurements on morphology of filopodia and intensity of IAPs staining are represented as box-plots. The significance for each condition on the measurements was tested using the non-parametric Mann-Whitney U test (Wilcoxon Rank-Sum test) for independent samples. All statistical analyses were performed using the R Stats Package version 3.3.1 (R Core Team, 2016).

## Figure legends

**Figure 3 – figure supplement 1.**
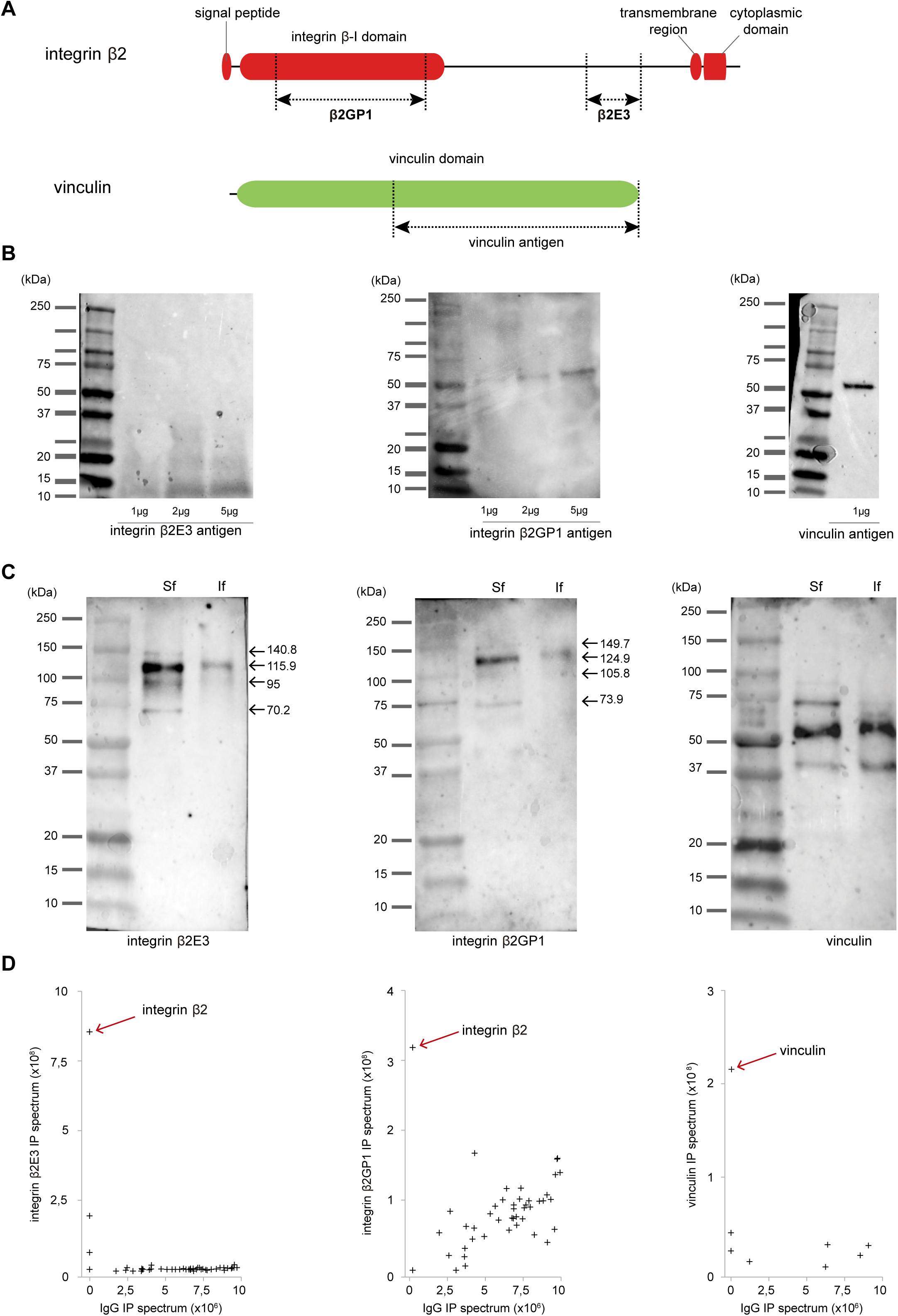
Integrin and vinculin antibodies validation by western-blot and mass-spectrometry. (A) Schematic of vinculin and integrin β2 proteins showing the predicted functional domains according to Pfam (El-Gebali et al. 2019), and the position of the protein fragments used as antigens for antibody production. (B) Western Blots of several concentrations of the purified antigens probed with integrin β2 and vinculin antibodies. (C) Western Blots of soluble (Sf) and insoluble (If) fractions of protein extracts from adherent *Capsaspora* cells probed with integrin and vinculin antibodies, show major bands at the predicted molecular weights. (D) Scatter plots showing the abundance of *Capsaspora* proteins detected by mass-spectrometry from immunoprecipitates obtained with integrin and vinculin antibodies relative to the IgG control, show the strong specificity of the antibodies.

**Figure 3 – figure supplement 2.**
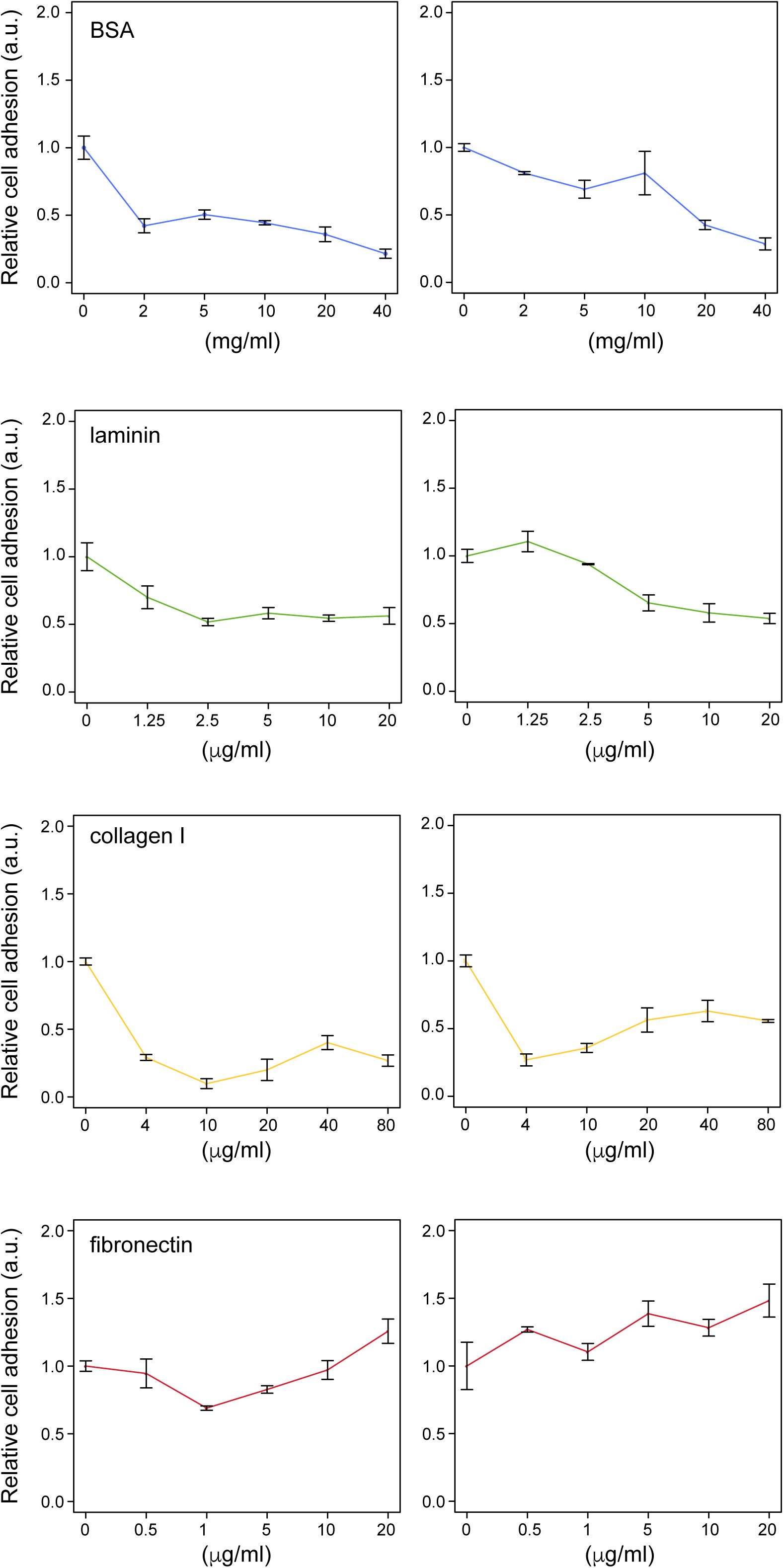
*Capsaspora owczarzaki* cell adhesion is enhanced on fibronectin-coated surfaces. Cell adhesion measurements on surfaces treated with increasing concentrations of BSA, laminin, collagen I and fibronectin. These experiments represent additional experiments compared to Figure 3. Relative cell adhesion was set as the ratio between the signal obtained between ligand-coated and uncoated surfaces. Data represents the mean ± s.e.m of 4 technical replicates.

Video 1. *Capsaspora owczarzaki* filopodia are used for locomotion.

Time-lapse of an adherent *Capsaspora* cell transfected with a membrane label (CoNMM:mcherry) (selected frames are shown in Figure 2D). Every frame corresponds to 10 sec.

